# Generalized neural decoders for transfer learning across participants and recording modalities

**DOI:** 10.1101/2020.10.30.362558

**Authors:** Steven M. Peterson, Zoe Steine-Hanson, Nathan Davis, Rajesh P. N. Rao, Bingni W. Brunton

## Abstract

**Objective:** Advances in neural decoding have enabled brain-computer interfaces to perform increasingly complex and clinically-relevant tasks. However, such decoders are often tailored to specific participants, days, and recording sites, limiting their practical long-term usage. Therefore, a fundamental challenge is to develop neural decoders that can robustly train on pooled, multi-participant data and generalize to new participants.

**Approach:** We introduce a new decoder, HTNet, which uses a convolutional neural network with two innovations: (1) a Hilbert transform that computes spectral power at data-driven frequencies and (2) a layer that projects electrode-level data onto predefined brain regions. The projection layer critically enables applications with intracranial electrocorticography (ECoG), where electrode locations are not standardized and vary widely across participants. We trained HTNet to decode arm movements using pooled ECoG data from 11 of 12 participants and tested performance on unseen ECoG or electroencephalography (EEG) participants; these pretrained models were also subsequently fine-tuned to each test participant.

**Main results:** HTNet outperformed state-of-the-art decoders when tested on unseen participants, even when a different recording modality was used. By fine-tuning these generalized HTNet decoders, we achieved performance approaching the best tailored decoders with as few as 50 ECoG or 20 EEG events. We were also able to interpret HTNet’s trained weights and demonstrate its ability to extract physiologically-relevant features.

**Significance:** By generalizing to new participants and recording modalities, robustly handling variations in electrode placement, and allowing participant-specific fine-tuning with minimal data, HTNet is applicable across a broader range of neural decoding applications compared to current state-of-the-art decoders.

## 1. Introduction

Brain-computer interfaces that interpret neural activity to control robotic or virtual devices have shown tremendous potential for assisting patients with neurological disabilities, including motor impairments, sensory deficits, and mood disorders [1–8]. At the same time, brain-computer interfaces offer new insights about the function of neural circuits, including how sensorimotor information is represented in the brain [9–11]. Advances in brain-computer interfaces have been driven in part by improved neural decoding algorithms [12, 13]. However, it can be difficult to collect enough data to train decoders, especially given the non-stationary nature of the recorded signals, leading to decoders that generalize poorly to new data and require frequent re-calibrations [14–17]. Alternatively, *generalized neural decoders* can be trained by pooling data across multiple participants [18–21]. Such generalized decoders must be robust to inter-participant differences and capable of fine-tuning with only a few training examples. By increasing decoder robustness and reducing the burden of repeated calibrations, generalized decoders have the potential to greatly enhance the practical long-term usage of brain-computer interfaces [22].

Frequency-domain techniques that extract spectral power features from time-domain recordings have long been shown to be useful in decoding neural population recordings. These techniques are especially well-suited to neural recordings such as intracranial electro-corticography (ECoG) and scalp electroencephalography (EEG) recordings, which contain oscillatory signals at specific frequency bands that correspond to different behaviors or neural phenomena [23–27]. In addition, relative spectral power patterns between a task and a baseline condition can be surprisingly similar across studies, even when measured with different neural recording modalities [28–33]. Such similarities motivate the use of spectral power features for generalized decoding. Because neural recordings are non-stationary, many decoders use instantaneous spectral power features, computed by band-pass filtering the data and then applying the Hilbert transform [17].

To make power spectral features directly comparable, inter-participant differences in electrode placement and frequency content must be addressed when developing generalized decoders. While EEG electrode coverage is typically standardized across participants, invasive ECoG electrode placement is clinically motivated and highly variable, making it difficult to align electrodes from one participant to the next [24, 34]. A similar cross-participant alignment issue occurs with EEG cortical dipoles following blind source separation [35]. To overcome these variable dipole locations, one successful approach has been to project EEG measures onto common brain regions using radial basis function interpolation [36, 37]. ECoG signals can similarly be projected from electrodes to common brain regions [38], but it has remained unclear how useful this method is for neural decoding. Even with aligned electrode placements, the frequency bands containing behaviorally-relevant spectral power can be highly variable across participants [38–40]. This variability is problematic when selecting band-pass filter cutoff frequencies prior to applying the Hilbert transform because researchers often rely on pre-existing knowledge of the neural signal, and traditional frequency bands may not apply to a particular task of interest. To decrease user bias, data-driven approaches have been proposed to analyze neural spectral power [41–43]. However, these techniques apply to frequency bands with distinct spectral power peaks and thus ignore other frequencies that might be useful for decoding. While a promising approach has been recently proposed [44], developing decoders that robustly handle variable frequency content remains an open problem.

We are motivated to address such cross-participant differences in frequency content by using convolutional neural networks, which generate data-driven features and can also be fine-tuned when presented with new data. Convolutional neural networks combine data-driven feature extraction with pattern recognition and have set the bar for state-of-the-art neural decoding performance for speech, motor imagery, and attention tasks [14, 45–49]. Importantly, trained convolutional neural networks can be fine-tuned to new data, a process made more efficient by freezing various network layers during re-training [50, 51]. Far from being “black-box” models, convolutional layers perform data-driven temporal and spatial filtering, and careful analysis of trained convolutional weights can be used to interpret the spatiotemporal features that are key for decoding [52–54]. Our approach builds on EEGNet—a compact convolutional neural network decoder that can be trained on small datasets, provides interpretable model structure, and has outperformed other deep learning decoders [55].

In this paper, we present *HTNet*, a convolutional neural network architecture that decodes neural data with variable electrode placements using data-driven spectral features projected onto common brain regions (Figure 1A). We developed HTNet by augmenting EEGNet with custom Hilbert transform and projection onto brain region layers. To characterize its decoding performance, we trained HTNet to classify naturalistic arm movement vs. rest using ECoG recordings from a dataset containing simultaneous video and ECoG recordings from 12 patients being monitored before neurosurgery in a hospital [38]. We then tested these trained decoders’ ability to generalize to unseen ECoG participants or to unseen EEG participants performing a similar behavioral task [56]. In both cases, HTNet was able to decode these naturalistic arm movements and consistently outperformed other state-of-the-art decoders. Furthermore, we fine-tuned pretrained HTNet decoders with a few events from individual participants, and the resultant fine-tuned decoders approach the performance of the best tailored decoders with only 50 ECoG or 20 EEG events. Finally, we interpreted HTNet’s trained weights to understand how it generalizes and found that it primarily relied on physiologically-relevant features at low-frequencies (<20 Hz) near the motor cortex. Our findings demonstrate that HTNet is a generalized decoder that robustly handles inter-participant and inter-modality variability and fine-tunes to new participants using minimal data. We believe our work advances the field of generalized neural decoders with an architecture that is robust, interpretable, and successful at generalizing across unseen participants and recording modalities.

**Figure 1:**
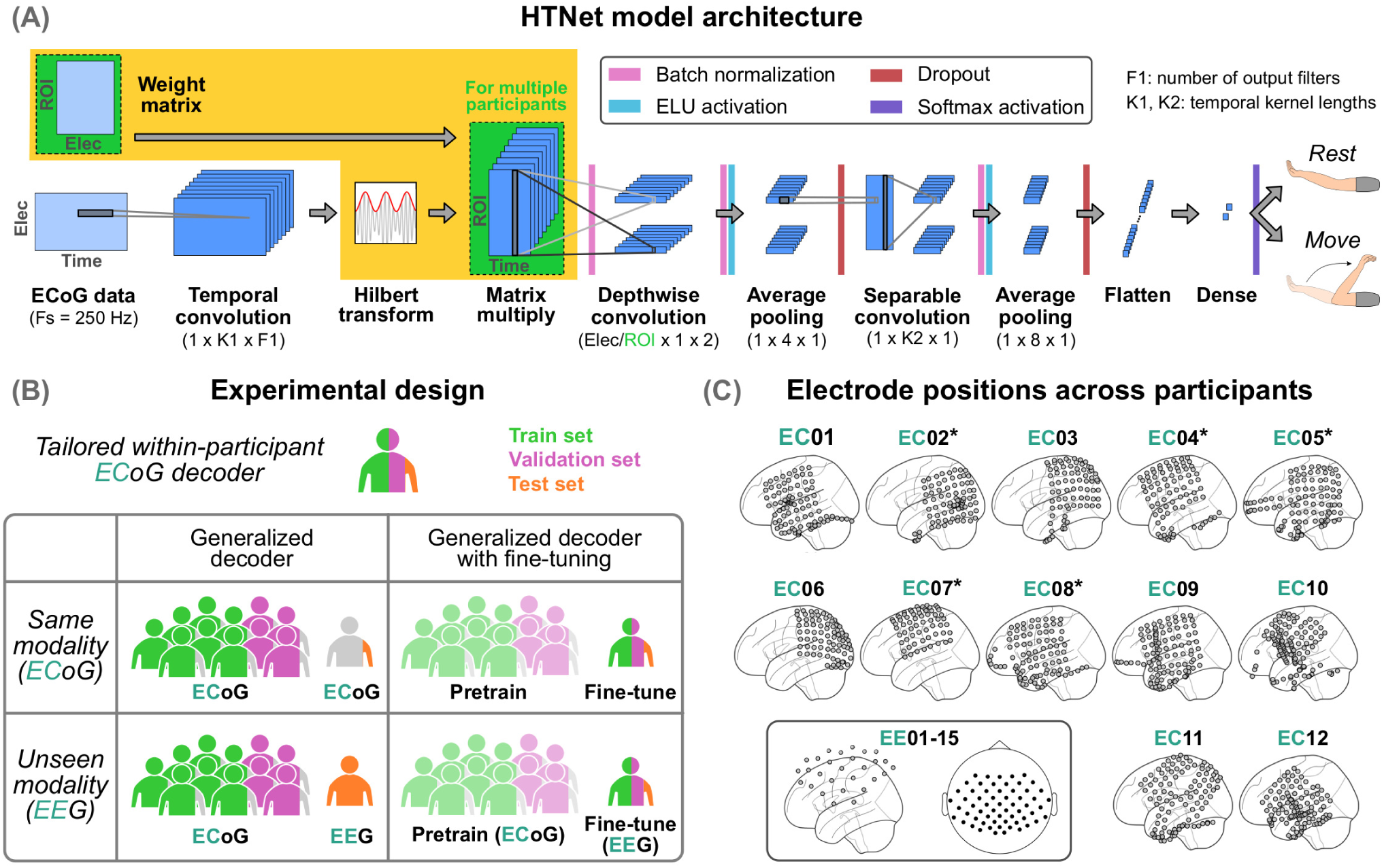
Overview of HTNet architecture, experimental design, and electrode locations. **(A)** HTNet is a convolutional neural network architecture that extends EEGNet [55] (differences shown in yellow) by handling cross-participant variations in electrode placement and frequency content. The temporal convolution and Hilbert transform layers generate data-driven spectral features that can then be projected from electrodes (Elec) onto common regions of interest (ROI) using a predefined weight matrix. **(B)** Using electrocorticography (ECoG) data, we trained both tailored within-participant and generalized multi-participant models to decode arm movement vs. rest. Multi-participant decoders were tested separately on held-out data from unseen participants recorded with either the same modality as the train set (ECoG) or an unseen modality (EEG). We then fine-tuned these pretrained decoders using data from the test participant. **(C)** Electrode placement varies widely among the 12 ECoG participants. Electrode coverage is sparser for the 15 EEG participants compared to ECoG, but both modalities overlap in coverage of sensorimotor cortices. Asterisks denote five participants whose electrodes were mirrored from the right hemisphere.

## 2. Methods

### 2.1. Intracranial electrocorticography (ECoG) dataset

We obtained concurrent ECoG and video recordings from 12 human participants (8 males, 4 females) during continuous clinical epilepsy monitoring conducted at Harborview Medical Center in Seattle, WA. These recordings lasted 7*±* 2 days per participant (mean *±* SD). Participants were aged 29 *±* 8 years old (mean *±* SD) and had electrodes implanted primarily in one hemisphere (5 right, 7 left). Our study was approved by the University of Washington Institutional Review Board for the protection of human participants, and all participants provided written informed consent. Our decoding task was to classify upper-limb “move” and “rest” events of the arm contralateral to the implanted electrode hemisphere. We obtained non-concurrent move and rest events from video recordings via markerless pose tracking and automated state segmentation (see Singh et al. [57] for further details). Move events correspond to wrist movement that occurred after at least 0.5 seconds of no movement, while rest events indicate no movement in either wrist for at least three seconds.

We performed ECoG data processing using custom MNE-Python scripts [58]. We first removed median DC drift and high-amplitude discontinuities. Each participant’s ECoG data was then band-pass filtered (1–200 Hz), notch filtered, and re-referenced to the common median across electrodes. We also removed noisy electrodes based on abnormal standard deviation (> 5 IQR) or kurtosis (> 10 IQR). Next, we generated 10-second ECoG segments centered around each “move” and “rest” event. ECoG segments with missing data or large artifacts were removed based on abnormal spectral power density. See Peterson et al. [38] for further ECoG pre-processing details. We then downsampled to 250 Hz and trimmed segments to two seconds centered around each event. For every participant, we balanced the number of move and rest segments within each recording day, resulting in 1155*±* 568 events per participant (mean *±* SD).

Electrode positions were localized using the Fieldtrip toolbox in Matlab [59, 60]. This process involved co-registering preoperative MRI and postoperative CT scans, manually selecting electrodes in 3D space, and warping electrode positions into Montreal Neurological Institute (MNI) space [61]. Using this common MNI coordinate system enabled us to directly compare electrode positions between ECoG participants (see Fig. 1C).

### 2.2. Comparative cross-modal dataset (EEG)

To test decoder generalizability across recording modalities, we used a publicly available EEG dataset of 15 human participants performing cued right elbow flexion movements [56]. Participants were aged 27 *±* 5 years old (mean *±* SD) and performed 60 movement and 60 rest trials each, resulting in 120 total events. Because only cue onset times were available, we determined the onset of each movement event by thresholding the hand’s radial displacement after it was cued to move.

EEG data was recorded at 512 Hz, notch filtered at 50 Hz, referenced to right mastoid, and band-pass filtered between 0.01–200 Hz. We pre-processed the data by average referencing, 1 Hz high-pass filtering, resampling to 250 Hz, and generating 2-second segments centered around each event. Each participant had 61 EEG electrodes, whose MNI positions were estimated using the 10-5 system template from Fieldtrip [60, 62].

### 2.3. Computing projection matrices

We accounted for variations in electrode placement by mapping electrode positions onto common brain regions based on distance as described below. To increase electrode overlap among ECoG participants, we mirrored all right hemisphere electrode positions onto the left hemisphere. Using EEGLAB and Matlab, we mapped from electrode positions to small, predefined brain regions by computing radial basis function kernel distances between each electrode and brain region (2 cm full-width at half-maximum) [36, 63, 64]. This projection procedure was performed separately for each participant. We projected to regions within sensorimotor areas, as defined by the AAL atlas [65] (precentral, postcentral, and inferior parietal), in order to limit the projected data size. We then normalized these distance values for every region so that each region’s values across all electrodes summed to one. These normalized distances created a projection matrix for each participant, which we later used to estimate the activity at each common region of interest by performing a weighted average of electrode-level data.

### 2.4. HTNet architecture

HTNet builds upon EEGNet [55], a compact convolutional neural network developed using Tensorflow and Keras. EEGNet has three convolution layers: (1) a one-dimensional convolution analogous to temporal band-pass filtering, (2) a depthwise convolution to perform spatial filtering, and (3) a separable convolution to identify temporal patterns across the previous filters. For HTNet, we added a Hilbert transform layer after this initial temporal convolution to compute relevant spectral power features using a data-driven filter-Hilbert analog (see Figure 1A). We then added a matrix multiplication layer to project electrode-level spectral power onto common brain regions of interest, using the pre-computed weight matrices described in the previous section. Note that the matrix multiplication layer was not necessary when the same participant was used for training and testing, as the electrodes remain consistent. All other HTNet layers were the same as EEGNet.

### 2.5. Data division and cross-validation

We compared HTNet decoding performance against EEGNet, random forest, and minimum distance decoders. The minimum distance decoder used Riemannian mean and distance values for classification [66,67]. We assessed decoder performance during three scenarios (Fig. 1B): (1) testing on an untrained recording day for the same ECoG participant (*tailored decoder*), (2) testing on an untrained ECoG participant (*same modality*), and (3) testing on participants from the EEG dataset after training only on the ECoG dataset (*unseen modality*). Note that we used the same trained, multi-participant ECoG decoders for same and unseen modality conditions. Additionally, we projected data onto common regions of interest for all decoders in order to enable reasonable decoding for the same and unseen modality conditions. For all scenarios, we performed 36 pseudo-random selections (folds) of the training and validation datasets, such that each of the 12 ECoG participants was the test participant three times. We used each ECoG participant’s last recording day as the test set (orange in Fig. 1B) and excluded it from all training and validation sets. All training, validation, and test sets were balanced with equal numbers of move and rest events. We used nonparametric statistics to test for significant effects of decoder type on test accuracy (Friedman test, *p* < 0.05) and significant pairwise differences among decoders (Wilcoxon signed-rank test with false discovery rate correction [68]).

### 2.6. Hyperparameter tuning

We performed hyperparameter tuning to identify optimal values for each decoder. We tuned HTNet and EEGNet simultaneously using six hyperparameters: temporal kernel length, separable kernel length, temporal filter count, dropout rate, dropout type, and model type (HTNet or EEGNet). For the random forest decoder, we tuned two hyperparameters: maximum depth and number of estimators. The minimum distance decoder had no tunable hyperparameters.

We tuned hyperparameters separately for the tailored decoder and same modality conditions, using the Optuna toolbox [69]. We ran 25 random forest parameter selections (trials) and 100 HTNet/EEGNet trials for each condition. Performance was measured using validation accuracy, averaged over 36 folds for tailored decoding or 12 folds for same modality. We sampled from parameter space using Optuna’s tree-structured Parzen estimator [70, 71], which selects optimal parameter values based on the performance during previous trials.

Overall, we found that hyperparameter selections minimally affected decoder performance (see Fig. S1). Still, we selected hyperparameter values from the trial with the highest validation accuracy (Table S1) for each condition to ensure optimal decoder performance.

### 2.7. Fine-tuning decoder performance to the test participant

In addition to testing generalizability, we assessed how much a generalized HTNet decoder improves when re-trained using data from the test participant, a process known as *fine-tuning*. Fine-tuning is a transfer learning technique where some layers of the pretrained model are “frozen” and not adjusted during re-training, reducing the number of parameters to fit [72]. We fine-tuned our pretrained same and unseen modality decoders using a portion of the test participant’s data. Additionally, we fine-tuned each HTNet convolutional layer separately and all layers together, resulting in four re-trained models per fold (see Fig. 3A). When separately tuning each convolutional layer, we also re-trained the nearby batch normalization layers, as shown in Fig. 3A, which notably boosted performance. We tested for significant differences among these four fine-tuning models using Wilcoxon signed-rank tests with false discovery rate correction.

**Figure 2:**
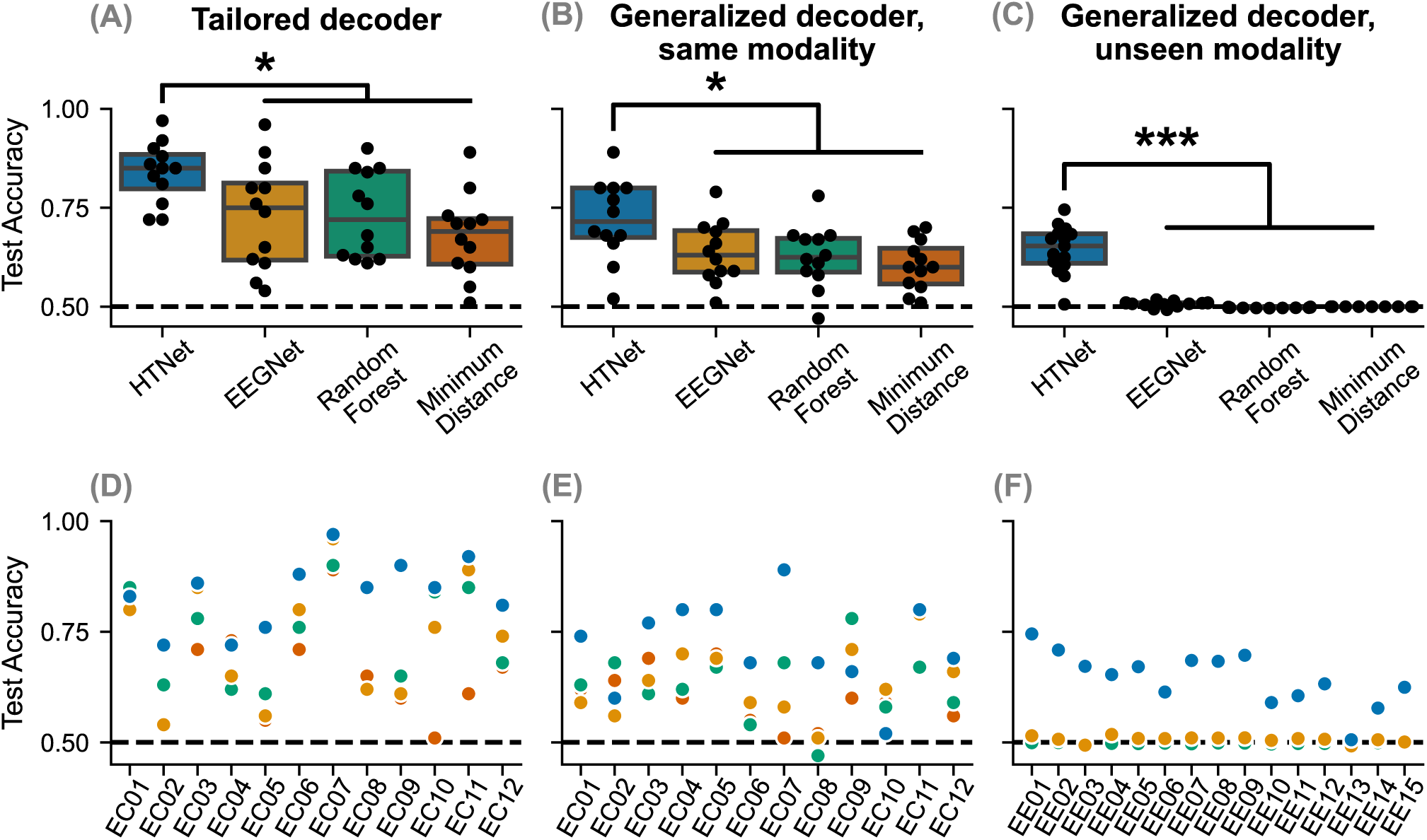
HTNet generalizes better than EEGNet and other decoders. HTNet achieves significantly higher test accuracy than EEGNet, random forest, and minimum distance decoders across all three scenarios: **(A)** tailored (*p* < 0.05), **(B)** same modality (*p* < 0.05), and **(C)** unseen modality (*p ≤* 0.001). Note that the trained models for same and unseen modality conditions are identical; only the test set differs. **(D–F)** Bottom row displays decoder performance grouped by test participant for each fold. For unseen modality, HTNet uses relative power to minimize cross-modal scaling differences, resulting in performance that is above chance (dashed line) despite only training on ECoG data.

**Figure 3:**
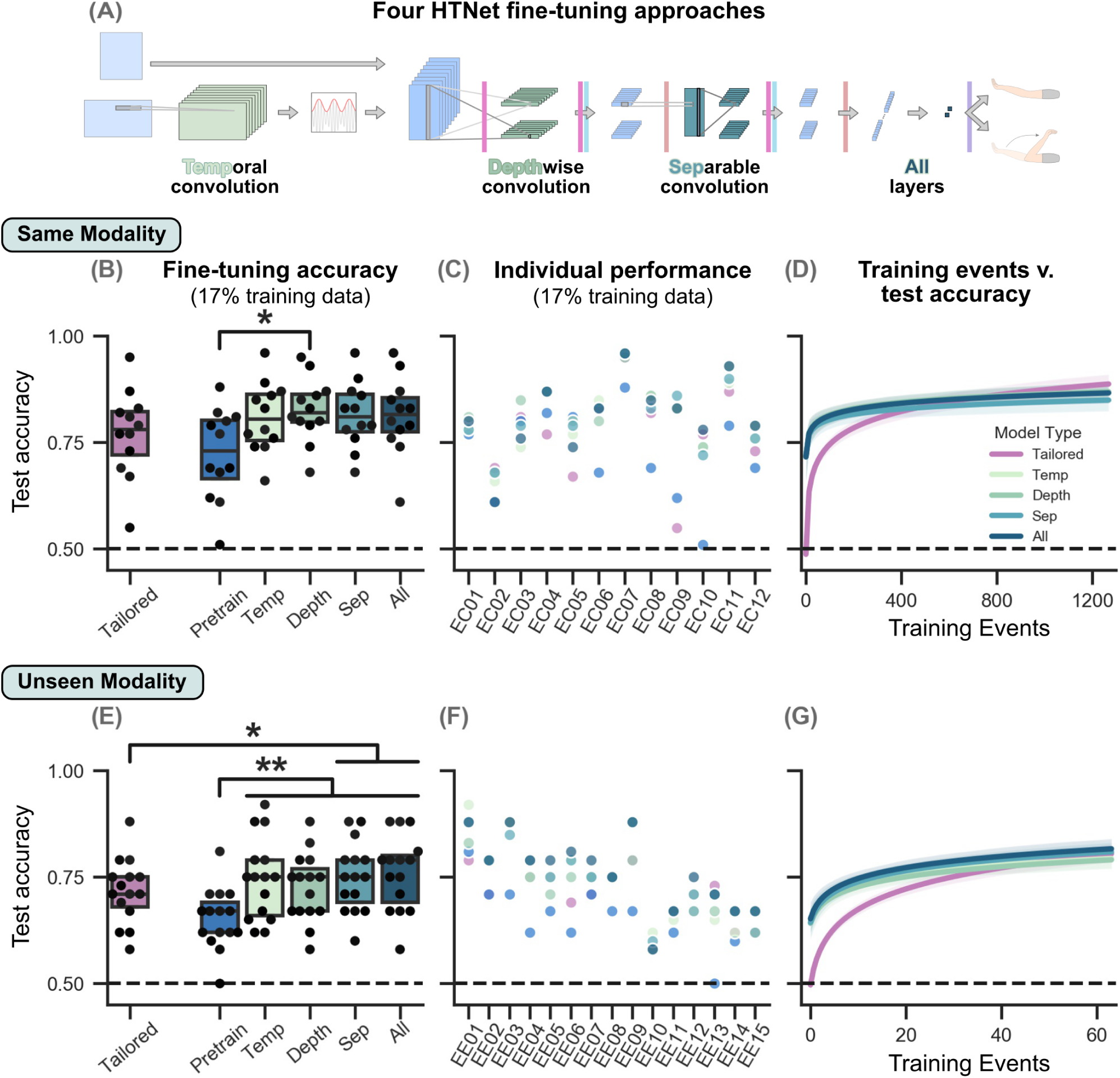
Fine-tuning HTNet improves performance, even when few training events are available. **(A)** For pretrained, multi-participant HTNet decoders, we separately fine-tuned each convolution layer (with nearby batch normalizations) and also fine-tuned all trainable layers. Layers with trainable parameters are highlighted. **(B, E)** For both same and unseen modality cases, we find no performance differences between fine-tuning approaches when training on 17% of available events, but our fine-tuned decoders often significantly improve test accuracy compared to the pretrained models and, in some cases, the randomly-initialized tailored decoders (* *p* < 0.05, ** *p ≤* 0.01). **(C, F)** Similarly, a breakdown of performance by participant shows that fine-tuning usually improves performance compared to the pretrained model (blue), but no single fine-tuning approach decodes consistently better than the others. **(D, G)** As we decreased the number of training events below 400 (same modality) or 30 (unseen modality), fine-tuned decoders generally achieved higher test accuracy than tailored decoders and remained close to accuracies achieved by tailored decoders trained on all available events. Lines show logarithmic fits for each group, with shading denoting the 95% confidence interval of the slope. Note that pretrained model accuracies are included for all fine-tuned models when there are 0 training events.

During fine-tuning, we also varied the amount of training/validation data available in order to assess its impact on fine-tuning performance. For all variations, the test set remained fixed as the last recording day for each ECoG participant and 30 randomly-selected events for every EEG participant. Using the remaining test participant data, we selected four amounts of training/validation data: 17% training/8% validation, 33% training/17% validation, 50% training/25% validation, and 67% training/33% validation. We then linearly modelled the relationship between test accuracy and the logarithm of the number of training events for each fine-tuning model. All training, validation, and test sets contained equal numbers of move and rest events. Note that the number of events used for fine-tuning differs across ECoG participants because the total number of events varies. In addition, we trained randomly-initialized HTNet decoders with the same training/validation data used for fine-tuning in order to compare tailored decoding with our fine-tuned models.

### 2.8. Comparing performance of HTNet spectral measures

Although we primarily used HTNet to generate data-driven spectral power features, HTNet can be easily adapted to generate other spectral measures that may boost decoding performance. These spectral measures still use the Hilbert transform, which can be used to find instantaneous power, phase, or frequency [17]. For the same three conditions, we tested four spectral measures: (1) log-transform of one plus power, (2) relative power, (3) unwrapped instantaneous phase, and (4) instantaneous frequency, also known as frequency sliding [73]. We tested for significant differences in test accuracy when HTNet decoders implemented one of these four measures or spectral power, using Wilcoxon signed-rank tests with false discovery rate correction.

When computing relative power, we first took the log-transform of one plus power and then subtracted out the average power from -1 to -0.5 seconds before each event. This procedure is analogous to baseline-subtraction of spectral power and should be robust to large-scale spectral power variations across days, participants, and recording modalities. Because of its potential robustness to large differences in signal scaling, relative power was also used in HTNet decoders that we used to compare with other decoder types, but only for the unseen modality condition.

### 2.9. Interpreting model weights

Like EEGNet, HTNet’s first two convolutional layers are interpretable and can indicate the spatiotemporal features used for decoding. To analyze the temporal features, we fed a white noise signal into HTNet’s trained temporal convolution and computed the frequency response magnitude using Welch’s method, similar to a Bode plot [74]. We then averaged frequency responses across temporal filters and folds. To determine important spatial features, we computed the absolute value of HTNet’s trained depthwise convolution weights [55], averaged across filters. This process generated one weight for each brain region of interest. We then scaled the maximum value per fold to one and averaged across folds.

While the temporal frequency response shows which frequencies were used for decoding, it does not show what the activity looks like at these frequencies. To visualize such activity, we computed the difference in log spectral power between move and rest conditions, projected onto a region near the motor cortex (Fig 4B far left region in the second row from the top). We took the difference between the average move and average rest log spectrograms for each participant and then averaged the resulting differences across participants for both the ECoG and EEG datasets.

**Figure 4:**
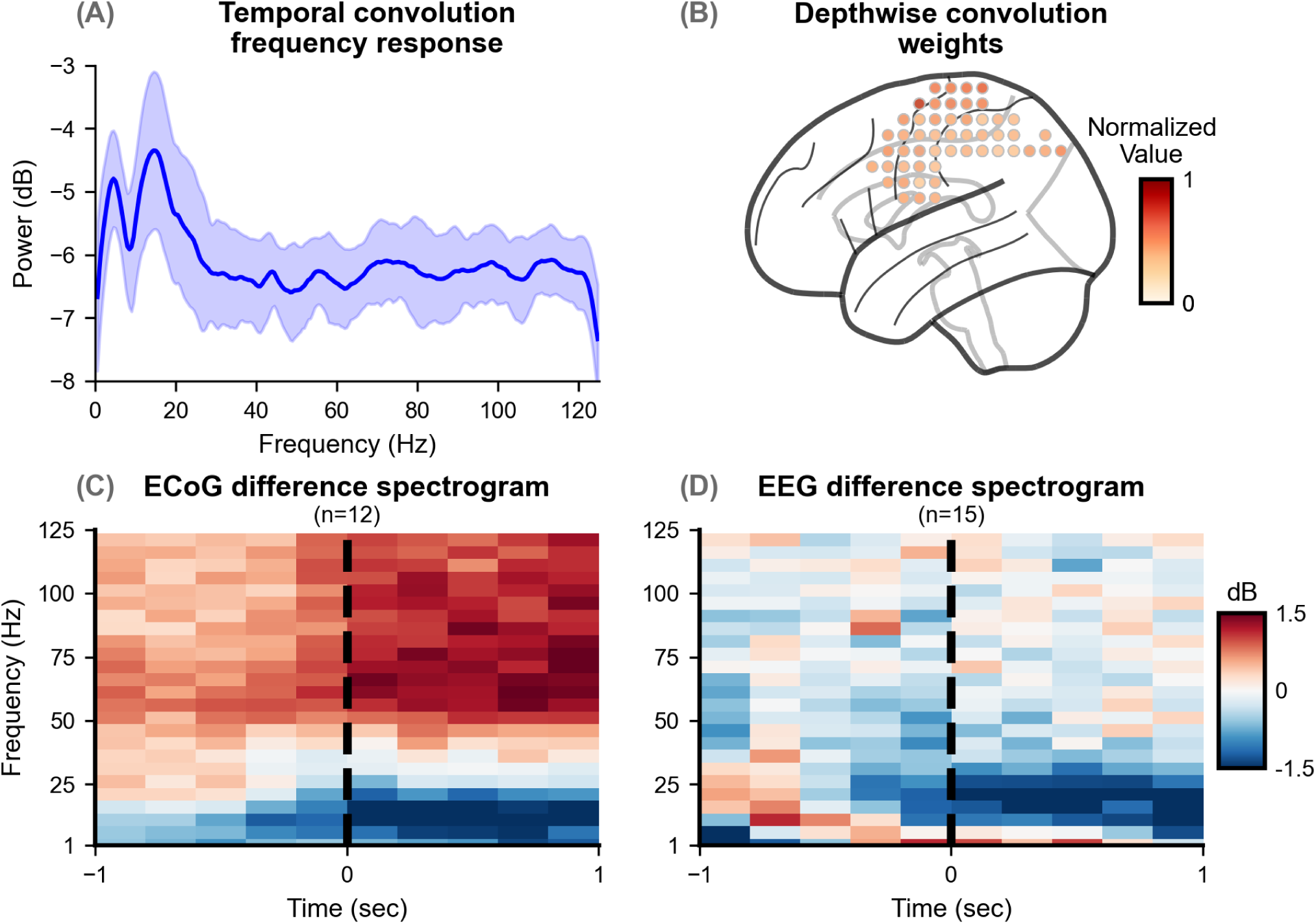
HTNet extracts physiologically-relevant features at low frequencies and near the motor cortex. By analyzing HTNet’s early convolution layers, we determined the types of spatial and temporal features consistently used for multi-participant decoding. **(A)** The temporal convolution layer’s frequency response, averaged across filters and folds, shows a consistent focus on low frequency features (<20 Hz). **(B)** Based on depthwise convolution weight magnitudes, cortical regions near the central sulcus and towards the midline were found to be consistently important for decoding, as expected for upper-limb movements. Out of the 144 total regions of interest used for decoding, we show here the 51 regions (shaded circles) located on the cortical surface. **(C–D)** Difference spectrograms between arm movement and rest events reveal a common low-frequency (<25 Hz) spectral power decrease near movement onset (0 sec) of similar magnitude across ECoG and EEG datasets.

### 2.10. Effect of training participants on performance

We also assessed how many training participants are needed for improved decoder performance. Increasing the number of training participants should improve decoder performance on an unseen test participant, but the improvement from adding new participants will likely diminish as more training participants are added. We varied the number of training participants from 1–10, always using one validation participant. Participants were pseudo-randomly selected across 36 folds such that each participant was the test participant three times. We linearly modelled the relationship between test accuracy and the logarithm of the number of training participants, identifying decoders with significant nonzero trends (*p* < 0.05, two-tailed t-test with false discovery rate correction).

### 2.11. Effect of electrode overlap on performance

Because electrode locations varied among ECoG participants, we also tested if higher decoding performance corresponded to increased electrode overlap between same modality training and test participants. We estimated electrode overlap between training and test participants using a custom fraction overlap metric that allowed us to combine multiple participants from the training set. For each sensorimotor region, we summed unnormalized projection matrix weights across electrodes to estimate how many electrodes were nearby. We then identified regions of high electrode coverage for each participant by thresholding summations greater than 0.07. Because these regions are common across participants, we could average these summation values across training participants prior to thresholding. Next, we divided the number of thresholded regions common to both training and test participants by the number of thresholded regions in the training set to obtain fraction overlap. A fraction overlap of 1.0 indicates that the test participant’s thresholded regions include all thresholded regions from the training participants. We linearly modelled the relationship between fraction overlap and test accuracy across folds and identified decoders with significant nonzero slope (*p* < 0.05, two-tailed t-test with false discovery rate correction [68]).

## 3 Results

Here, we show that our approach to movement decoding, HTNet, is generalizable and tunable, learning common patterns from the training data that transfer to unseen participants and recording modalities. Through a series of systematic experiments, we use decoders tailored to each participant—by far the most common approach to training decoders—as a standard to which the performance of our generalized decoders are compared (Figure 1B). In particular, our training data are from 12 ECoG participants during uninstructed, naturalistic arm movements (Figure 1C, [38, 57]); our test data are then either one ECoG participant withheld from the training set, or participants from an entirely independent EEG dataset. HTNet consistently out-performed other decoders, and fine-tuning pre-trained HTNet decoders with a small number of the unseen participant’s events yielded decoders that approached the performance of tailored decoders trained on many more events. Further, we show that HTNet works by extracting physiologically-relevant spectral features from the data.

### 3.1. Decoder generalization

HTNet consistently outperformed other decoders in all conditions, including tailored within-participant decoding, generalized decoding to unseen participants in the same modality, and generalized decoding to unseen modality participants (Figure 2). For each condition, we found a significant effect of decoder type on test accuracy (tailored: *p* = 3.94e − 4, same modality: *p* = 0.034, unseen modality: *p* = 1.95e − 8 respectively; Friedman test). For tailored decoding, HTNet achieved test accuracy of 84% ± 8% (mean ±SD), which was significantly higher than EEGNet (73% ± 14%, *p* = 0.003), random forest (73% ± 11%, *p* = 0.004), and minimum distance decoders (68% ± 11%, *p* = 0.010; Wilcoxon signed-rank test with false discovery rate correction). For same modality, HTNet test accuracy of 72% ± 10% was again significantly increased compared to EEGNet (64%±8%, *p* = 0.042), random forest (63% ± 8%, *p* = 0.037), and minimum distance decoders (60% ± 6%, *p* = 0.021). No other significant pairwise differences were found for either condition.

In the unseen modality condition, HTNet was the only decoder to generalize above chance. All pairwise comparisons were statistically significant (*p* < 0.05) in the unseen modality condition, due to low variability in cross-participant test accuracy for each non-HTNet decoder. However, HTNet’s unseen modality test accuracy was, on average, *∼*15% higher than all other decoders and the only decoder to perform well-above chance (50%). Therefore, we only show the pairwise statistical results in Figure 2C between HTNet and each of the other three decoders. Average computational time during training is shown for each decoder type in Tables S2 and S3. In addition to spectral power, we also developed HTNet models that decoded using instantaneous phase and frequency features (Fig. S2 and Tables S4, S5). Briefly, HTNet with phase performed worse than spectral power for tailored and same modality conditions, while using instantaneous frequency resulted in similar performance to HTNet with spectral power.

In addition to outperforming the other decoders on average, we show in Figure 2D–F that HTNet was the single best arm movement decoder for almost every individual participant in all conditions. HTNet performance was also consistently well-above random chance for all but two participants (EC10 same modality and EE13), much more than any other decoder. HTNet’s consistently high accuracy demonstrates its ability to robustly generalize to a variety of participants with differences in electrode placement and signal quality.

### 3.2. Fine-tuning generalized decoders

By fine-tuning these pretrained HTNet decoders using as few as 50 events from the held-out participant, HTNet can approach performances of the best tailored decoders trained using all available participant data. We tested fine-tuning each convolutional layer separately as well as re-training all layers of the network, as shown in Fig. 3A. We found no significant differences in performance between our fine-tuning approaches, but we did find significant improvements in test accuracy when comparing fine-tuned decoders to pretrained decoders or tailored decoders trained on the same data, even when training on only 17% of the test participant’s available events. For same modality (Fig. 3B), fine-tuning just the depthwise convolution significantly increased test accuracy compared to the pretrained models (*p* = 0.049; Wilcoxon signed-rank test with false discovery rate correction). In the unseen modality condition (Fig. 3E), all fine-tuning approaches significantly increased test accuracy compared to the pretrained models (temporal convolution: *p* = 0.0054, depthwise convolution: *p* = 0.0094, separable convolution: *p* = 0.0054, all layers: *p* = 0.0054). Additionally, fine-tuning the separable convolution and all HTNet layers resulted in significantly higher test accuracy than tailored HTNet decoders (*p* = 0.021 and *p* = 0.015, respectively). No other comparisons were statistically significant.

Similarly, decoder performance for each participant shows that, in general, fine-tuning HTNet increases test accuracy. All tested fine-tuning approaches yielded similar performance (Fig. 3C,F). In addition, computation times are consistent across fine-tuning approaches (Tables S6, S7). By varying the number of events used (Fig. 3D,G), we find that fine-tuning approaches outperform randomly-initialized, tailored decoders when fewer than *∼*400 ECoG or *∼*30 EEG events from the test participant are available for training (see Fig. S3 for separated plots of each fine-tuning approach). In addition, fine-tuned HTNet decoders approach performances of the best tailored decoders with as few as 50 ECoG or 20 EEG events available for training. Finally, we demonstrated that we can fine-tune HTNet decoders to ECoG events after pretraining on EEG events (Fig. S4).

### 3.3. Interpreting network computations

Trained HTNet models achieve generalized decoding by extracting physiologically-relevant features, specifically at low frequencies and near the motor cortex (Fig. 4). HTNet’s temporal and depthwise convolutional layers are interpretable, so we can probe these trained layers to determine the spatiotemporal features most often used for decoding, just like with EEG-Net [55]. Here, we analyzed the trained layers of generalized HTNet decoders that were used for the same and unseen modality conditions.

Across decoders, we find a consistent emphasis on low-frequency (<20 Hz) temporal features (Fig. 4A) when analyzing the average frequency response over temporal convolution filters. These decoders also frequently focused on cortical regions near the motor cortex (towards the central sulcus and midline), based on trained depthwise convolution weights (Fig. 4B). When we take the difference in spectral power between arm movement and rest events near these motor cortical regions, we find, consistent with previous ECoG and EEG studies [75, 76], that low-frequency spectral power decreases during movement onset, with a similar magnitude for both ECoG and EEG data. This consistent magnitude in low-frequency power between recording modalities likely explains why our trained HTNet models generalized to the EEG participants prior to any fine-tuning.

Another crucial factor in HTNet performance is the spatial overlap of electrodes between the training and test participants, as well as how many participants were used for training (Fig. 5). We observe significant positive trends between electrode fraction overlap and test accuracy for HTNet (0.33 slope, *p* = 0.005; t-test with false discovery rate correction) and EEGNet (0.23, *p* = 0.025). We also find significant logarithmic relationships between the number of training participants and test accuracy for HTNet (0.07 slope, *p* = 2.77e − 7), EEGNet (0.06, *p* = 2.77e − 7, and random forest (0.03, *p* = 0.003). We initially tried a linear fit, but found that logarithmic scaling of the number of training events resulted in a better fit. This logarithmic trend suggests that, at a certain point, adding more participants will not noticeably improve decoder performance. The number of training participants also affects computation time and number of epochs used during training (Fig. S5). Our findings suggest that HTNet is best able to incorporate information when many training participants are used, especially those with similar electrode placement to the test participant.

**Figure 5:**
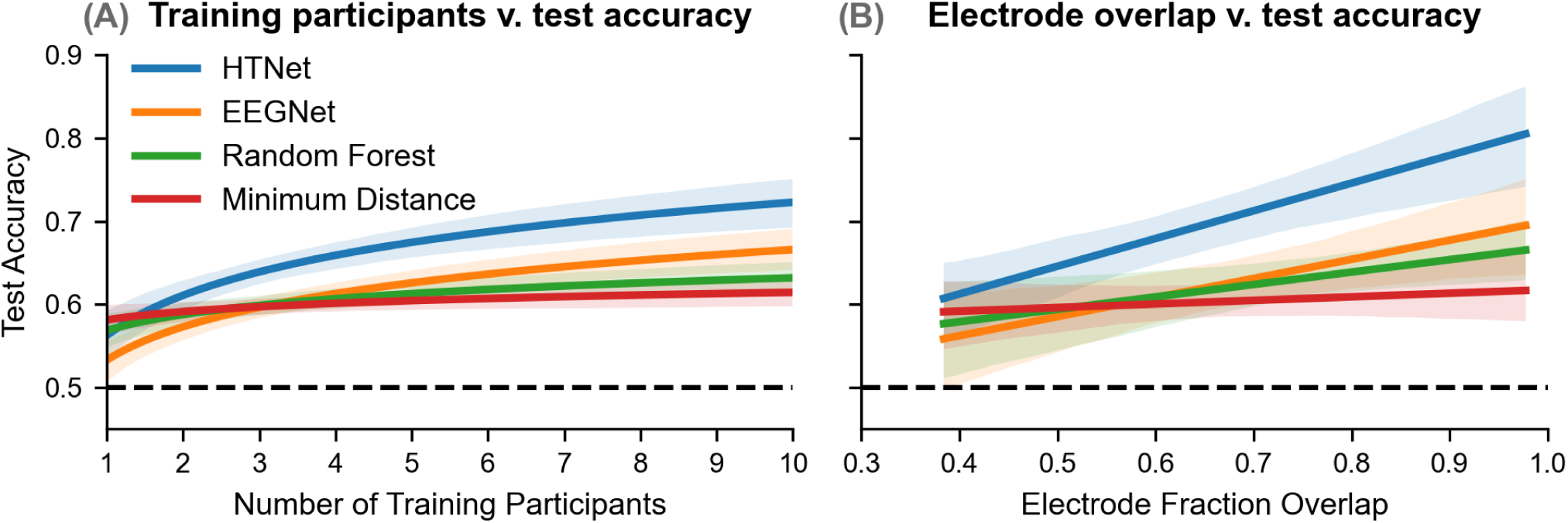
HTNet performance improves with increases in training participants and electrode overlap. **(A)** Adding more training participants significantly improved test accuracy for HTNet, EEGNet, and random forest decoders (*p* < 0.05). The line of best fit is shown for each group, with shading denoting the 95% confidence interval of the slope. The logarithmic relationship seen suggests that performance will not substantially improve once a certain number of training participants are used. **(B)** We also compared test accuracy as a function of electrode coverage overlap between the training and test participants and found a significant positive relationship only for HTNet and EEGNet (*p* < 0.05).

## 4. Discussion

We demonstrated that HTNet can outperform state-of-the-art neural decoders when generalizing to new participants, even when a different recording modality is used. Developed as an extension of EEGNet [55], our key contributions to the neural network architecture are the addition of a Hilbert transform layer and a weight matrix to project individual electrode locations onto common brain regions. These trained HTNet decoders can be fine-tuned to a new test participant with fewer than 100 of the test participant’s events and still decode almost as well as tailored decoders that have been trained on substantially more events. To achieve this performance, HTNet decoders consistently extract physiologically-relevant features, as revealed by our analysis of trained decoder weights.

To our knowledge, HTNet is the first decoder that can generalize and transfer its learning across both ECoG participants and different recording modalities. Previous studies have implemented decoders that can transfer across different EEG devices [14, 46, 77, 78] or leverage data from concurrent recording modalities [79, 80], but none of these decoders have demonstrated the ability to generalize to an entirely different recording modality. As for generalizing across participants, many decoders can do this with EEG data [14], including EEGNet [55], but development of analogous ECoG decoders has been hindered by the high variation in electrode placement across ECoG patients. Similarly, several studies have implemented fine-tuning when decoding EEG or ECoG data [21, 50, 81–84], but these decoders were trained on either one participant or non-brain data, instead of multiple ECoG participants. By training on multiple participants’ data, HTNet can generalize effectively to unseen participants and avoid overfitting to any one participant.

When we compared performance across different types of decoders, HTNet consistently outperformed EEGNet, even in the tailored condition. These performance differences between HTNet and EEGNet arise solely from HTNet’s Hilbert transform layer because for all decoders, we projected onto common brain regions when training on multiple participants. We used the Hilbert transform to compute spectral power, which unlike the time-domain signal does not include a phase component. When we used just this phase component for decoding, we found that HTNet performance substantially worsened compared to using spectral power (see Fig. S2). This difference in accuracy suggests that decoding directly from the time-domain signal is suboptimal for our specific decoding task because the phase component is less informative than spectral power. Based on this insight, we would expect EEGNet to perform as well as or better than HTNet when decoding neural data with highly informative phase, such as event-related potentials [85, 86], depending on whether HTNet uses phase or spectral power to decode.

More broadly, HTNet demonstrates the value of integrating computational models, such as deep learning, with insights from neural signal processing. Fusing computational methods with scientific insight has inspired novel solutions that leverage the strengths of both approaches [45, 53, 87]. An alternative to our approach would have been to develop an end-to-end neural network model that simply learns to compute spectral power or project electrode-level signals onto a common grid. However, incorporating explicit transformations to spectral power and common cortical regions minimized the number of trainable parameters (and hence the amount of training data needed) and, like EEGNet, kept HTNet’s initial layers interpretable. Furthermore, we were able to apply insights from previous ECoG/EEG research by explicitly baseline-subtracting spectral power within HTNet in order to generalize from ECoG to unseen EEG data [88, 89]. On the other hand, deep learning models can generate data-driven features that may be computationally expensive to obtain using other available methods. While many data-driven spatial filtering methods are available [17, 90], identifying frequencies with relevant spectral power often requires either a brute force search or applying techniques such as wavelet convolution to compute power at several frequencies, increasing the size of an already high-dimensional dataset [91, 92]. In contrast, HTNet converges quickly and provides a low-dimensional feature representation in the spectral domain. We believe that further embedding of neural signal processing into data-driven methods such as deep learning will continue to enhance the robustness and generalizability of future neural decoders.

Our study has several important limitations to consider. First, we classified two event types (arm movement versus rest), which is substantially less than the types of complex behaviors present in many neural decoding paradigms. Nonetheless, HTNet’s architecture allows for decoding more than two types of events, and EEGNet has been shown to perform well when decoding among four types of behavior [55]. We chose ECoG and EEG datasets from a task with only two event types because the electrode positions were in a common coordinate system (MNI), which was essential when projecting data onto common brain regions for multi-participant decoding. Our analysis also may have been limited by projecting only onto 144 sensorimotor brain regions, which we chose based on the decoding task. For real-world decoders, the most informative regions may not be clear, requiring data-driven region-selection approaches to avoid high memory usage and slow computation times [37, 38]. Finally, we could have constrained HTNet’s temporal convolution layer to learn more meaningful narrow-band filters [93].

We are currently exploring extensions of HTNet for a variety of applications such as cross-frequency coupling [94, 95], long-term state decoding [6], cross-task decoding [96], and data-driven regression [97, 98]. In addition, other decoding measures could be substituted for the Hilbert transform, including non-Fourier methods [99, 100], and more complex interpolation schemes could be used to generate the projection matrix by incorporating participant-specific cortical anatomy [101, 102]. Besides ECoG and EEG, HTNet may also be useful for generalizing across participants with stereotactic EEG or local field potential recordings [103, 104]. Overall, HTNet provides a useful decoding framework applicable across a variety of tasks and overcomes important obstacles towards developing robust, generalized neural decoders that can be fine-tuned with minimal data.

### Code and data availability

Our HTNet code is publicly available at: https://github.com/BruntonUWBio/HTNet_generalized_decoding. The code in this repository can be used in conjunction with publicly available ECoG (https://figshare.com/projects/Generalized_neural_decoders_for_transfer_learning_across_participants_and_recording_modalities/90287) and EEG [56] datasets to generate all of the main findings and figures from our study.

## Acknowledgements

We thank Satpreet Singh, Ellie Strandquist, Nancy Wang, Pierre Karashchuk, and Kurt Weaver for discussions and help with data analysis and model design, along with neurosurgeons Jeffrey G. Ojemann, Andrew Ko, and the excellent staff at Harborview Hospital Neurosurgery department for their care of the patients during their monitoring. This research was supported by funding from the Defense Advanced Research Projects Agency (FA8750-18-2-0259), the National Science Foundation (1630178 and EEC-1028725), and the Washington Research Foundation.

## Notes

https://figshare.com/projects/Generalized_neural_decoders_for_transfer_learning_across_participants_and_recording_modalities/90287

https://github.com/BruntonUWBio/HTNet_generalized_decoding

### Competing Interest Statement

The authors have declared no competing interest.

## References

[1] Patrick D. Ganzer, Samuel C. Colachis, Michael A. Schwemmer, David A. Friedenberg, Collin F. Dunlap, Carly E. Swiftney, Adam F. Jacobowitz, Doug J. Weber, Marcia A. Bockbrader, and Gaurav Sharma. Restoring the sense of touch using a sensorimotor demultiplexing neural interface. Cell, 181(4):763–773.e12, 2020.

[2] Kai J Miller, Dora Hermes, and Nathan P Staff. The current state of electrocorticography-based brain– computer interfaces. Neurosurgical Focus, 49(1):E2, 2020.

[3] Ksenia Volkova, Mikhail A Lebedev, Alexander Kaplan, and Alexei Ossadtchi. Decoding movement from electrocorticographic activity: A review. Frontiers in neuroinformatics, 13, 2019.

[4] Soroush Niketeghad and Nader Pouratian. Brain machine interfaces for vision restoration: The current state of cortical visual prosthetics. Neurotherapeutics, 16(1):134–143, 2019.

[5] Stephanie Martin, Josédel R. Millán, Robert T. Knight, and Brian N. Pasley. The use of intracranial recordings to decode human language: Challenges and opportunities. Brain and Language, 193:73–83, 2019.

[6] Omid G Sani, Yuxiao Yang, Morgan B Lee, Heather E Dawes, Edward F Chang, and Maryam M Shanechi. Mood variations decoded from multi-site intracranial human brain activity. Nature Biotechnology, 36(10):954–961, 2018.

[7] Wei Wang, Jennifer L. Collinger, Alan D. Degenhart, Elizabeth C. Tyler-Kabara, Andrew B. Schwartz, Daniel W. Moran, Douglas J. Weber, Brian Wodlinger, Ramana Vinjamuri, Robin C. Ashmore, John W. Kelly, and Michael L. Boninger. An electrocorticographic brain interface in an individual with tetraplegia. PLoS ONE, 8, 2013.

[8] Eric C. Leuthardt, Gerwin Schalk, Jonathan R. Wolpaw, Jeffrey G. Ojemann, and Daniel W. Moran. A brain-computer interface using electrocorticographic signals in humans. Journal of neural engineering, 1 2:63–71, 2004.

[9] Alan D Degenhart, William E Bishop, Emily R Oby, Elizabeth C Tyler-Kabara, Steven M Chase, Aaron P Batista, and M Yu Byron. Stabilization of a brain–computer interface via the alignment of low-dimensional spaces of neural activity. Nature Biomedical Engineering, pages 1–14, 2020.

[10] Emily R Oby, Jay A Hennig, Aaron P Batista, M Yu Byron, and Steven M Chase. Intracortical brain– machine interfaces. In Neural Engineering, pages 185–221. Springer, 2020.

[11] Jennifer L Collinger, Robert A Gaunt, and Andrew B Schwartz. Progress towards restoring upper limb movement and sensation through intracortical brain-computer interfaces. Current Opinion in Biomedical Engineering, 8:84–92, 2018.

[12] Xiaotong Gu, Zehong Cao, Alireza Jolfaei, Peng Xu, Dongrui Wu, Tzyy-Ping Jung, and Chin-Teng Lin. Eeg-based brain-computer interfaces (bcis): A survey of recent studies on signal sensing technologies and computational intelligence approaches and their applications. arXiv preprint 2001.11337, 2020.

[13] Rajesh P. N. Rao. Brain-Computer Interfacing: An Introduction. Cambridge University Press, Cambridge, 2013.

[14] Dongrui Wu, Yifan Xu, and Bao-Liang Lu. Transfer learning for eeg-based brain-computer interfaces: A review of progress made since 2016, 2020.

[15] Jan Van Erp, Fabien Lotte, and Michael Tangermann. Brain-computer interfaces: beyond medical applications. Computer, 45(4):26–34, 2012.

[16] Gan Huang, Guangquan Liu, Jianjun Meng, Dingguo Zhang, and Xiangyang Zhu. Model based generalization analysis of common spatial pattern in brain computer interfaces. Cognitive neurodynamics, 4(3):217–223, 2010.

[17] Mike X Cohen. Analyzing Neural Time Series Data: Theory and Practice, jan 2014.

[18] Chuanqi Tan, Fuchun Sun, Tao Kong, Wenchang Zhang, Chao Yang, and Chunfang Liu. A survey on deep transfer learning. In ICANN, 2018.

[19] Martin Volker, Robin T. Schirrmeister, Lukas D. J. Fiederer, Wolfram Burgard, and Tonio Ball. Deep transfer learning for error decoding from non-invaahive EEG. In 2018 6th International Conference on Brain-Computer Interface (BCI), pages 1–6. IEEE, 2018.

[20] Ivan Zubarev, Rasmus Zetter, Hanna-Leena Halme, and Lauri Parkkonen. Adaptive neural network classifier for decoding meg signals. Neuroimage, 197:425–434, 2019.

[21] Ahmed M. Azab, Lyudmila Mihaylova, Kai Keng Ang, and Mahnaz Arvaneh. Weighted transfer learning for improving motor imagery-based brain–computer interface. IEEE Transactions on Neural Systems and Rehabilitation Engineering, 27(7):1352–1359, 2019.

[22] Clemens Brunner, Niels Birbaumer, Benjamin Blankertz, Christoph Guger, Andrea Kübler, Donatella Mattia, José del R Millán, Felip Miralles, Anton Nijholt, Eloy Opisso, et al. Bnci horizon 2020: towards a roadmap for the bci community. Brain-computer interfaces, 2(1):1–10, 2015.

[23] Kai J. Miller. A library of human electrocorticographic data and analyses. Nature Human Behaviour, 3(11):1225–1235, 2019.

[24] Josef Parvizi and Sabine Kastner. Promises and limitations of human intracranial electroencephalography. Nature Neuroscience, 21(4):474–483, 2018.

[25] Kana Takaura, Naotsugu Tsuchiya, and Naotaka Fujii. Frequency-dependent spatiotemporal profiles of visual responses recorded with subdural ecog electrodes in awake monkeys: Differences between high- and low-frequency activity. NeuroImage, 124:557–572, 2016.

[26] Aysegul Gunduz, Peter Brunner, Amy Daitch, Eric C Leuthardt, Anthony L Ritaccio, Bijan Pesaran, and Gerwin Schalk. Neural correlates of visual-spatial attention in electrocorticographic signals in humans. Frontiers in human neuroscience, 5:89–89, 09 2011.

[27] Tobias Pistohl, Tonio Ball, Andreas Schulze-Bonhage, Ad Aertsen, and Carsten Mehring. Prediction of arm movement trajectories from ecog-recordings in humans. Journal of Neuroscience Methods, 167(1):105–114, 2008.

[28] Anne B Martin, Xiaofang Yang, Yuri B Saalmann, Liang Wang, Avgusta Shestyuk, Jack J Lin, Josef Parvizi, Robert T Knight, and Sabine Kastner. Temporal dynamics and response modulation across the human visual system in a spatial attention task: An ecog study. Journal of Neuroscience, 39(2):333–352, 2019.

[29] Steven M Peterson and Daniel P Ferris. Differentiation in theta and beta electrocortical activity between visual and physical perturbations to walking and standing balance. eneuro, 5(4), 2018.

[30] Franz Hell, Paul CJ Taylor, Jan H Mehrkens, and Kai Bötzel. Subthalamic stimulation, oscillatory activity and connectivity reveal functional role of stn and network mechanisms during decision making under conflict. Neuroimage, 171:222–233, 2018.

[31] Jun Jiang, Kira Bailey, and Xiao Xiao. Midfrontal theta and posterior parietal alpha band oscillations support conflict resolution in a masked affective priming task. Frontiers in human neuroscience, 12:175, 2018.

[32] Baltazar Zavala, Huiling Tan, Keyoumars Ashkan, Thomas Foltynie, Patricia Limousin, Ludvic Zrinzo, Kareem Zaghloul, and Peter Brown. Human subthalamic nucleus–medial frontal cortex theta phase coherence is involved in conflict and error related cortical monitoring. Neuroimage, 137:178–187, 2016.

[33] Takako Fujioka, Laurel J Trainor, Edward W Large, and Bernhard Ross. Internalized timing of isochronous sounds is represented in neuromagnetic beta oscillations. Journal of Neuroscience, 32(5):1791–1802, 2012.

[34] G. Schalk and E. C. Leuthardt. Brain-computer interfaces using electrocorticographic signals. IEEE Reviews in Biomedical Engineering, 4:140–154, 2011.

[35] Julie Onton, Marissa Westerfield, Jeanne Townsend, and Scott Makeig. Imaging human eeg dynamics using independent component analysis. Neuroscience & biobehavioral reviews, 30(6):808–822, 2006.

[36] Nima Bigdely-Shamlo, Tim Mullen, Kenneth Kreutz-Delgado, and Scott Makeig. Measure projection analysis: a probabilistic approach to eeg source comparison and multi-subject inference. NeuroImage, 72:287–303, 05 2013.

[37] Steven M Peterson, Estefania Rios, and Daniel P. Ferris. Transient visual perturbations boost short-term balance learning in virtual reality by modulating electrocortical activity. Journal of neurophysiology, 120 4:1998–2010, 2018.

[38] Steven M. Peterson, Satpreet H. Singh, Nancy X. R. Wang, Rajesh P. N. Rao, and Bingni W. Brunton. Behavioral and neural variability of naturalistic arm movements. bioRxiv, 2020.

[39] Simanto Saha and Mathias Baumert. Intra-and inter-subject variability in eeg-based sensorimotor brain computer interface: a review. Frontiers in Computational Neuroscience, 13:87, 2019.

[40] Tanja Krumpe, Katrin Baumgaertner, Wolfgang Rosenstiel, and Martin Spüler. Non-stationarity and inter-subject variability of eeg characteristics in the context of bci development. In GBCIC, 2017.

[41] Andrew W Corcoran, Phillip M Alday, Matthias Schle-sewsky, and Ina Bornkessel-Schlesewsky. Toward a reliable, automated method of individual alpha frequency (iaf) quantification. Psychophysiology, 55(7):e13064, 2018.

[42] Asrul Adam, Mohd Ibrahim Shapiai, Mohd Zaidi Mohd Tumari, Mohd Saberi Mohamad, and Marizan Mubin. Feature selection and classifier parameters estimation for eeg signals peak detection using particle swarm optimization. The Scientific World Journal, 2014, 2014.

[43] AKI Chiang, CJ Rennie, PA Robinson, JA Roberts, MK Rigozzi, RW Whitehouse, RJ Hamilton, and E Gordon. Automated characterization of multiple alpha peaks in multi-site electroencephalograms. Journal of Neuroscience Methods, 168(2):396–411, 2008.

[44] Michael X Cohen. A data-driven method to identify frequency boundaries in multichannel electrophysiology data. bioRxiv, 2020.

[45] Gopala K. Anumanchipalli, Josh Chartier, and Edward F. Chang. Speech synthesis from neural decoding of spoken sentences. Nature, 568(7753):493–498, 2019.

[46] Lichao Xu, Minpeng Xu, Yufeng Ke, Xingwei An, Shuang Liu, and Dong Ming. Cross-dataset variability problem in eeg decoding with deep learning. Frontiers in Human Neuroscience, 14, 2020.

[47] Yurui Ming, Weiping Ding, Danilo Pelusi, Dongrui Wu, Yu-Kai Wang, Mukesh Prasad, and Chin-Teng Lin. Subject adaptation network for eeg data analysis. Applied Soft Computing, 84:105689, 2019.

[48] Yannick Roy, Hubert Banville, Isabela Albuquerque, Alexandre Gramfort, Tiago H Falk, and Jocelyn Faubert. Deep learning-based electroencephalography analysis: a systematic review. Journal of neural engineering, 16(5):051001, 2019.

[49] Xiang Zhang, Lina Yao, Xianzhi Wang, Jessica Monaghan, David Mcalpine, and Yu Zhang. A survey on deep learning based brain computer interface: Recent advances and new frontiers. arXiv preprint 1905.04149, 2019.

[50] Joseph G. Makin, David A. Moses, and Edward F. Chang. Machine translation of cortical activity to text with an encoder–decoder framework. Nature Neuroscience, 23(4):575–582, 2020.

[51] Joos Behncke, Robin Tibor Schirrmeister, Martin Volker, Jiri Hammer, Petr Marusic, Andreas Schulze-Bonhage, Wolfram Burgard, and Tonio Ball. Cross-paradigm pre-training of convolutional networks improves intracranial eeg decoding. In 2018 IEEE International Conference on Systems, Man, and Cybernetics (SMC), pages 1046–1053. IEEE, 2018.

[52] Siavash Sakhavi, Cuntai Guan, and Shuicheng Yan. Learning temporal information for brain-computer interface using convolutional neural networks. IEEE transactions on neural networks and learning systems, 29(11):5619–5629, 2018.

[53] Robin Tibor Schirrmeister, Jost Tobias Springenberg, Lukas Dominique Josef Fiederer, Martin Glasstetter, Katharina Eggensperger, Michael Tangermann, Frank Hutter, Wolfram Burgard, and Tonio Ball. Deep learning with convolutional neural networks for eeg decoding and visualization. Human brain mapping, 38(11):5391–5420, 2017.

[54] Pouya Bashivan, Irina Rish, Mohammed Yeasin, and Noel Codella. Learning representations from eeg with deep recurrent-convolutional neural networks. arXiv preprint arXiv:1511.06448, 2015.

[55] Vernon J Lawhern, Amelia J Solon, Nicholas R Waytowich, Stephen M Gordon, Chou P Hung, and Brent J Lance. Eegnet: a compact convolutional neural network for eeg-based brain–computer interfaces. Journal of Neural Engineering, 15(5):056013, 2018.

[56] Patrick Ofner, Andreas Schwarz, Joana Pereira, and Gernot R. Müller-Putz. Upper limb movements can be decoded from the time-domain of low-frequency eeg. PLOS ONE, 12(8):1–24, 08 2017.

[57] Satpreet H. Singh, Steven M. Peterson, Rajesh P. N. Rao, and Bingni W. Brunton. Towards naturalistic human neuroscience and neuroengineering: behavior mining in long-term video and neural recordings, 2020.

[58] Alexandre Gramfort, Martin Luessi, Eric Larson, Denis Engemann, Daniel Strohmeier, Christian Brodbeck, Roman Goj, Mainak Jas, Teon Brooks, Lauri Parkkonen, and Matti Hamäläinen. Meg and eeg data analysis with mne-python. Frontiers in Neuroscience, 7:267, 2013.

[59] Arjen Stolk, Sandon Griffin, Roemer van der Meij, Callum Dewar, Ignacio Saez, Jack J. Lin, Giovanni Piantoni, Jan-Mathijs Schoffelen, Robert T. Knight, and Robert Oostenveld. Integrated analysis of anatomical and electrophysiological human intracranial data. Nature Protocols, 13(7):1699–1723, 2018.

[60] Robert Oostenveld, Pascal Fries, Eric Maris, and Jan-Mathijs Schoffelen. Fieldtrip: Open source software for advanced analysis of meg, eeg, and invasive electrophysiological data. Computational intelligence and neuroscience, 2011:156869–156869, 2011.

[61] Vladimir Fonov, Alan C. Evans, Kelly Botteron, C. Robert Almli, Robert C. McKinstry, and D. Louis Collins. Unbiased average age-appropriate atlases for pediatric studies. NeuroImage, 54(1):313–327, 2011.

[62] Robert Oostenveld and Peter Praamstra. The five percent electrode system for high-resolution eeg and erp measurements. Clinical Neurophysiology, 112(4):713–719, 2001.

[63] Jean-Philippe Vert, Koji Tsuda, and Bernhard Schölkopf. A primer on kernel methods. Kernel methods in computational biology, 47:35–70, 2004.

[64] Arnaud Delorme and Scott Makeig. Eeglab: an open source toolbox for analysis of single-trial eeg dynamics including independent component analysis. Journal of Neuroscience Methods, 134(1):9 –21, 2004.

[65] N. Tzourio-Mazoyer, B. Landeau, D. Papathanassiou, F. Crivello, O. Etard, N. Delcroix, B. Mazoyer, and M. Joliot. Automated anatomical labeling of activations in spm using a macroscopic anatomical parcellation of the mni mri single-subject brain. NeuroImage, 15(1):273–289, 2002.

[66] Marco Congedo, Alexandre Barachant, and Rajendra Bhatia. Riemannian geometry for eeg-based brain-computer interfaces; a primer and a review. Brain-Computer Interfaces, 4(3):155–174, 2017.

[67] Florian Yger, Maxime Berar, and Fabien Lotte. Riemannian approaches in brain-computer interfaces: a review. IEEE Transactions on Neural Systems and Rehabilitation Engineering, 25(10):1753–1762, 2016.

[68] Yoav Benjamini and Yosef Hochberg. Controlling the false discovery rate: A practical and powerful approach to multiple testing. Journal of the Royal Statistical Society. Series B (Methodological), 57(1):289–300, 1995.

[69] Takuya Akiba, Shotaro Sano, Toshihiko Yanase, Takeru Ohta, and Masanori Koyama. Optuna: A next-generation hyperparameter optimization framework. In Proceedings of the 25rd ACM SIGKDD International Conference on Knowledge Discovery and Data Mining, 2019.

[70] James Bergstra, Daniel Yamins, and David Cox. Making a science of model search: Hyperparameter optimization in hundreds of dimensions for vision architectures. In International conference on machine learning, pages 115–123, 2013.

[71] James S Bergstra, Rémi Bardenet, Yoshua Bengio, and Balázs Kégl. Algorithms for hyper-parameter optimization. In Advances in neural information processing systems, pages 2546–2554, 2011.

[72] Jason Yosinski, Jeff Clune, Yoshua Bengio, and Hod Lipson. How transferable are features in deep neural networks? In Advances in neural information processing systems, pages 3320–3328, 2014.

[73] Michael X Cohen. Fluctuations in oscillation frequency control spike timing and coordinate neural networks. The Journal of Neuroscience, 34(27):8988, 07 2014.

[74] RK Rao Yarlagadda. Analog and digital signals and systems, volume 1. Springer, 2010.

[75] Jae W Chung, Edward Ofori, Gaurav Misra, Christopher W Hess, and David E Vaillancourt. Beta-band activity and connectivity in sensorimotor and parietal cortex are important for accurate motor performance. NeuroImage, 144:164–173, 2017.

[76] Kai J. Miller, Eric C. Leuthardt, Gerwin Schalk, Rajesh P.N. Rao, Nicholas R. Anderson, Daniel W. Moran, John W. Miller, and Jeffrey G. Ojemann. Spectral changes in cortical surface potentials during motor movement. Journal of Neuroscience, 27(9):2424–2432, 2007.

[77] Dongrui Wu, Vernon J. Lawhern, W. David Hairston, and Brent J. Lance. Switching EEG headsets made easy: Reducing offline calibration effort using active weighted adaptation regularization. IEEE Transactions on Neural Systems and Rehabilitation Engineering, 24(11):1125–1137, 2016.

[78] Masaki Nakanishi, Yu-Te Wang, Chun-Shu Wei, Kuan-Jung Chiang, and Tzyy-Ping Jung. Facilitating calibration in high-speed BCI spellers via leveraging cross-device shared latent responses. IEEE Transactions on Biomedical Engineering, 67(4):1105–1113, 2020.

[79] Jordan Muraskin, Truman R Brown, Jennifer M Walz, Tao Tu, Bryan Conroy, Robin I Goldman, and Paul Sajda. A multimodal encoding model applied to imaging decision-related neural cascades in the human brain. NeuroImage, 180:211–222, 2018.

[80] Sarwat Fatima and Awais M Kamboh. Decoding brain cognitive activity across subjects using multimodal m/eeg neuroimaging. In 2017 39th Annual International Conference of the IEEE Engineering in Medicine and Biology Society (EMBC), pages 3224–3227. IEEE, 2017.

[81] Venkatesh Elango, Aashish N Patel, Kai J Miller, and Vikash Gilja. Sequence transfer learning for neural decoding. bioRxiv, 2017.

[82] Ran Wang, Xupeng Chen, Amirhossein Khalilian-Gourtani, Zhaoxi Chen, Leyao Yu, Adeen Flinker, and Yao Wang. Stimulus speech decoding from human cortex with generative adversarial network transfer learning. In 2020 IEEE 17th International Symposium on Biomedical Imaging (ISBI), pages 390–394. IEEE, 2020.

[83] Sharanya Arcot Desai, Thomas Tcheng, and Martha Morrell. Transfer-learning for differentiating epileptic patients who respond to treatment based on chronic ambulatory ECoG data. In 2019 9th International IEEE/EMBS Conference on Neural Engineering (NER), pages 1–4. IEEE, 2019.

[84] Axel Uran, Coert Van Gemeren, Rosanne van Diepen, Ricardo Chavarriaga, and José del R Millán. Applying transfer learning to deep learned models for eeg analysis. arXiv preprint arXiv:1907.01332, 2019.

[85] Ruslan Aydarkhanov Aydarkhanov, Marija Uscumlic, Ricardo Chavarriaga, Lucian Gheorghe, and Jose del R Millan. Spatial covariance improves bci performance for late erps components with high temporal variability. Journal of Neural Engineering, 2020.

[86] Emily S Kappenman and Steven J Luck. Best practices for event-related potential research in clinical populations. Biological psychiatry: cognitive neuroscience and neuroimaging, 1(2):110–115, 2016.

[87] Ziqian Xie, Odelia Schwartz, and Abhishek Prasad. Decoding of finger trajectory from ecog using deep learning. Journal of neural engineering, 15(3):036009, 2018.

[88] Li Hu, P Xiao, ZG Zhang, André Mouraux, and Gian Domenico Iannetti. Single-trial time–frequency analysis of electrocortical signals: Baseline correction and beyond. Neuroimage, 84:876–887, 2014.

[89] Lucas C Parra, Clay D Spence, Adam D Gerson, and Paul Sajda. Recipes for the linear analysis of eeg. Neuroimage, 28(2):326–341, 2005.

[90] Scott Makeig, Anthony J Bell, Tzyy-Ping Jung, and Terrence J Sejnowski. Independent component analysis of electroencephalographic data. In Advances in neural information processing systems, pages 145–151, 1996.

[91] Hyeon Kyu Lee and Young-Seok Choi. Application of continuous wavelet transform and convolutional neural network in decoding motor imagery brain-computer interface. Entropy, 21(12):1199, 2019.

[92] Zied Tayeb, Juri Fedjaev, Nejla Ghaboosi, Christoph Richter, Lukas Everding, Xingwei Qu, Yingyu Wu, Gordon Cheng, and Jörg Conradt. Validating deep neural networks for online decoding of motor imagery movements from eeg signals. Sensors, 19(1):210, 2019.

[93] Mirco Ravanelli and Yoshua Bengio. Interpretable convolutional filters with sincnet. ArXiv, abs/1811.09725, 2018.

[94] Roemer van der Meij, Michael Kahana, and Eric Maris. Phase–amplitude coupling in human electrocorticography is spatially distributed and phase diverse. Journal of Neuroscience, 32(1):111–123, 2012.

[95] Ryan T Canolty and Robert T Knight. The functional role of cross-frequency coupling. Trends in cognitive sciences, 14(11):506–515, 2010.

[96] He He and Dongrui Wu. Different set domain adaptation for brain-computer interfaces: A label alignment approach. IEEE Transactions on Neural Systems and Rehabilitation Engineering, 28(5):1091–1108, 2020.

[97] Yuqi Cui, Yifan Xu, and Dongrui Wu. Eeg-based driver drowsiness estimation using feature weighted episodic training. IEEE transactions on neural systems and rehabilitation engineering, 27(11):2263–2273, 2019.

[98] Dongrui Wu, Brent J Lance, Vernon J Lawhern, Stephen Gordon, Tzyy-Ping Jung, and Chin-Teng Lin. Eeg-based user reaction time estimation using riemannian geometry features. IEEE Transactions on Neural Systems and Rehabilitation Engineering, 25(11):2157–2168, 2017.

[99] Scott Cole and Bradley Voytek. Cycle-by-cycle analysis of neural oscillations. Journal of neurophysiology, 122(2):849–861, 2019.

[100] Scott R Cole and Bradley Voytek. Brain oscillations and the importance of waveform shape. Trends in cognitive sciences, 21(2):137–149, 2017.

[101] LLW Owen, TA Muntianu, AC Heusser, PM Daly, KW Scangos, and JR Manning. A gaussian process model of human electrocorticographic data. Cerebral Cortex (New York, NY: 1991), 2020.

[102] M Vermaas, MC Piastra, TF Oostendorp, NF Ramsey, and PHE Tiesinga. Femfuns: A volume conduction modeling pipeline that includes resistive, capacitive or dispersive tissue and electrodes. Neuroinformatics, 2020.

[103] Christian Herff, Dean J Krusienski, and Pieter Kubben. The potential of stereotactic-eeg for brain-computer interfaces: Current progress and future directions. Frontiers in Neuroscience, 14:123, 2020.

[104] Andrew Jackson and Thomas M Hall. Decoding local field potentials for neural interfaces. IEEE Transactions on Neural Systems and Rehabilitation Engineering, 25(10):1705–1714, 2016.

