## Supplementary Information for "Generalized neural decoders for transfer learning across participants and recording modalities"

| Hyperparameter | **Tailored Decoder** | **Same Modality** |
|---|---|---|
| Dropout Rate | 0.69 | 0.34 |
| Temporal Kernel Length (K1) | 64 | 24 |
| Temporal Filter Count (F1) | 20 | 19 |
| Separable Kernel Length (K2) | 56 | 88 |
| Dropout Type | Spatial Dropout 2D | Dropout |
| Number of Estimators | 240 | 240 |
| Maximum Depth | 9 | 6 |

**Table S1: Optimal parameter values from hyperparameter tuning.** Parameter values are shown for the hyperparameter tuning run with the highest accuracy on the validation data. We tuned hyperparameters separately for tailored decoder and same modality conditions. The first five parameters are for HTNet and EEGNet decoders, while the last two are for random forest decoders; the minimum distance decoder we used had no trainable parameters. Note that we used the same exact trained decoders for the same and unseen modality conditions; only the test set differed.

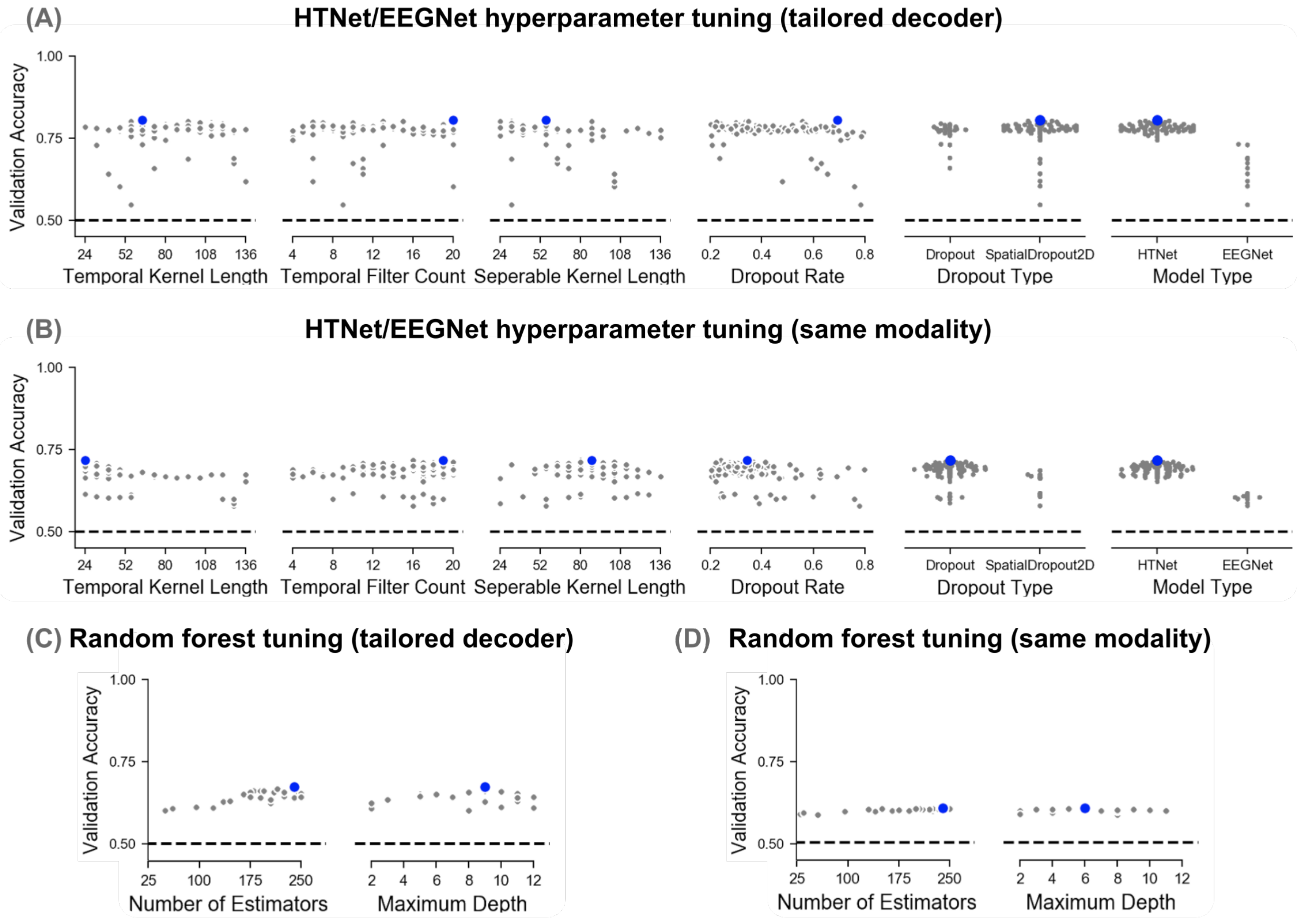


**Figure S1: Most hyperparameter selections do not greatly affect decoder performance.** For each condition, we performed 100 selections of six hyperparameters for HTNet/EEGNet decoders **(A–B)** and 25 selections of 2 hyperparameters for random forest **(C–D)**. We estimated performance using accuracy on a validation dataset, averaged over the 36 data folds. Blue dots indicate the parameter values for the trial with the highest validation accuracy in each condition. In general, our hyperparameter selections do not substantially alter decoder performance, except when selecting model type (HTNet vs. EEGNet).

| | HTNet | EEGNet | Random Forest | Minimum Distance |
|---|---|---|---|---|
| Tailored Decoder | 117.7 (89.4) | 66.4 (49.8) | 12.2 (8.0) | 6.9 (5.7) |
| Same Modality | 430.3 (173.1) | 185.0 (63.6) | 99.6 (12.8) | 247.7 (33.2) |

**Table S2: Training times across decoder types.** The average training time, in seconds, to train each decoder is shown with standard deviation in parentheses. While HTNet takes that longest time to finish training in both conditions, it still takes only a few minutes to train.

| | HTNet | EEGNet |
|---|---|---|
| Tailored Decoder | 55.0 (32.5) | 72.1 (46.9) |
| Same Modality | 9.6 (8.8) | 6.6 (8.3) |

**Table S3: Number of epochs during training for neural network decoders.** The average number of training epochs are shown for HTNet and EEGNet decoders with standard deviation in parentheses. Because both HTNet and EEGNet had early stopping criteria during training, the number of epochs used during training varied across runs. In general, both decoders trained over a similar number of epochs. Interestingly, same modality decoders needed fewer than 10 training epochs on average, far fewer than the number of training epochs during tailored decoding.

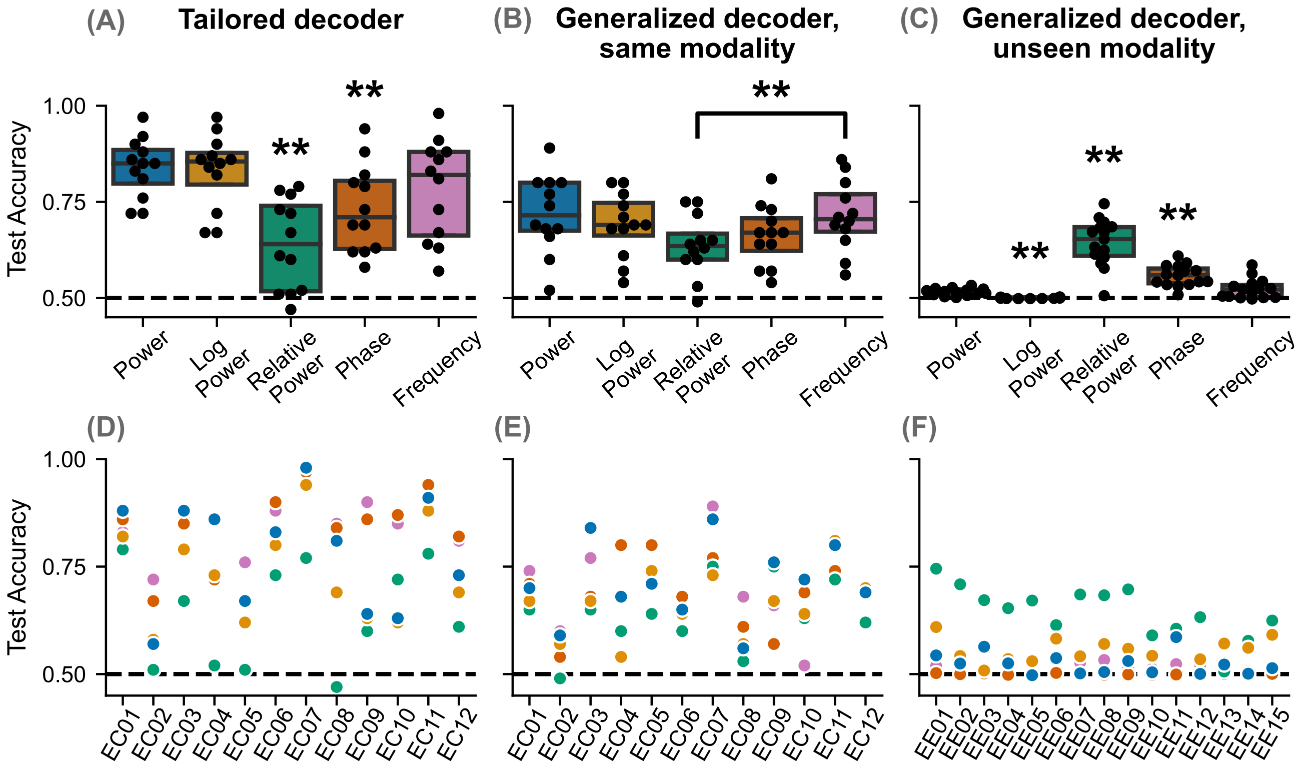


**Figure S2: HTNet can compute a variety of spectral measures.** HTNet's Hilbert transform layer allows us to compute instantaneous power, phase, and frequency measures that can be used for decoding. During tailored decoding, relative power and phase performed significantly worse than all other measures used ($p < 0.01$ for all; Wilcoxon signed-rank test with false discovery rate correction). For same modality, we only found a significant difference between instantaneous frequency and relative power. In the unseen modality condition, relative power, phase, and log power each performed significantly different from the other measures ($p < 0.01$), with relative phase having the highest test accuracy. Our findings indicate that relative power and phase are sub-optimal measures for decoding within our ECoG dataset, but they generalize well across recording modalities.

| | Power | Log Power | Relative Power | Phase | Frequency |
|---|---|---|---|---|---|
| Tailored Decoder | 117.7 (89.4) | 112.8 (77.3) | 69.9 (42.1) | 485.0 (458.7) | 549.6 (508.3) |
| Same Modality | 430.3 (173.1) | 431.0 (199.9) | 367.3 (54.6) | 1393.1 (601.9) | 1311.7 (450.9) |

**Table S4: HTNet training times when different spectral measures are used.** The average training time, in seconds, to train HTNet is shown when different power, phase, and frequency measures are used (standard deviation in parentheses). While all spectral power measures result in similar training times, instantaneous phase and frequency take approximately four times as long to train.

| | Power | Log Power | Relative Power | Phase | Frequency |
|---|---|---|---|---|---|
| Tailored Decoder | 55.0 (32.5) | 46.8 (24.8) | 17.4 (10.2) | 55.0 (43.2) | 57.5 (37.7) |
| Same Modality | 9.6 (8.8) | 8.0 (6.1) | 8.0 (4.2) | 16.1 (15.7) | 12.8 (11.3) |

**Table S5: Number of epochs when training HTNet with different spectral measures.** The average number of epochs needed to train HTNet decoders are shown when various spectral measures are used (standard deviation in parentheses). For all measures except relative power, tailored decoding needs about 5–6 times as many training epochs as same modality decoding. The tailored relative power decoders used one third as many training epochs as the decoders with other spectral measures and might have substantially benefited from having more time to train.

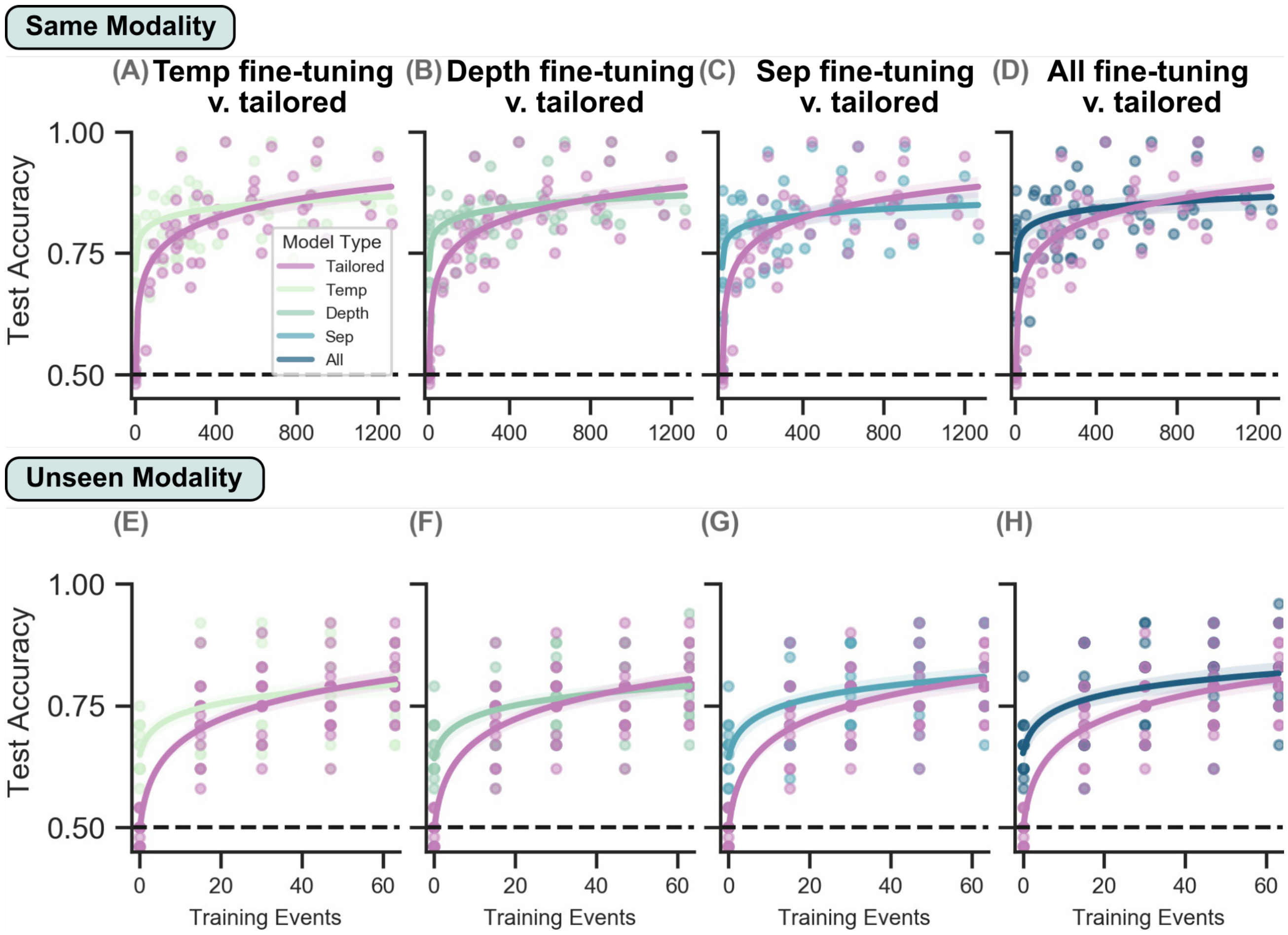


**Figure S3: HTNet decoder performance as the number of training events varies, separated by fine-tuning approach.** The relationship between test accuracy and the number of events used for fine-tuning is shown separately for each of the four fine-tuning approaches: **(A, E)** temporal convolution (Temp), **(B, F)** depthwise convolution (Depth), **(C, G)** separable convolution (Sep), and **(D, H)** all trainable layers (All). Each fine-tuning approach is compared to the performance of a randomly-initialized, tailored decoder trained on the same test participant data. Logarithmic lines of best fit are shown for each decoder type, with shading indicating the 95% confidence interval of the slope. Dots denote the single fold test accuracy when trained on either 17%, 33%, 50%, or 67% of available data.

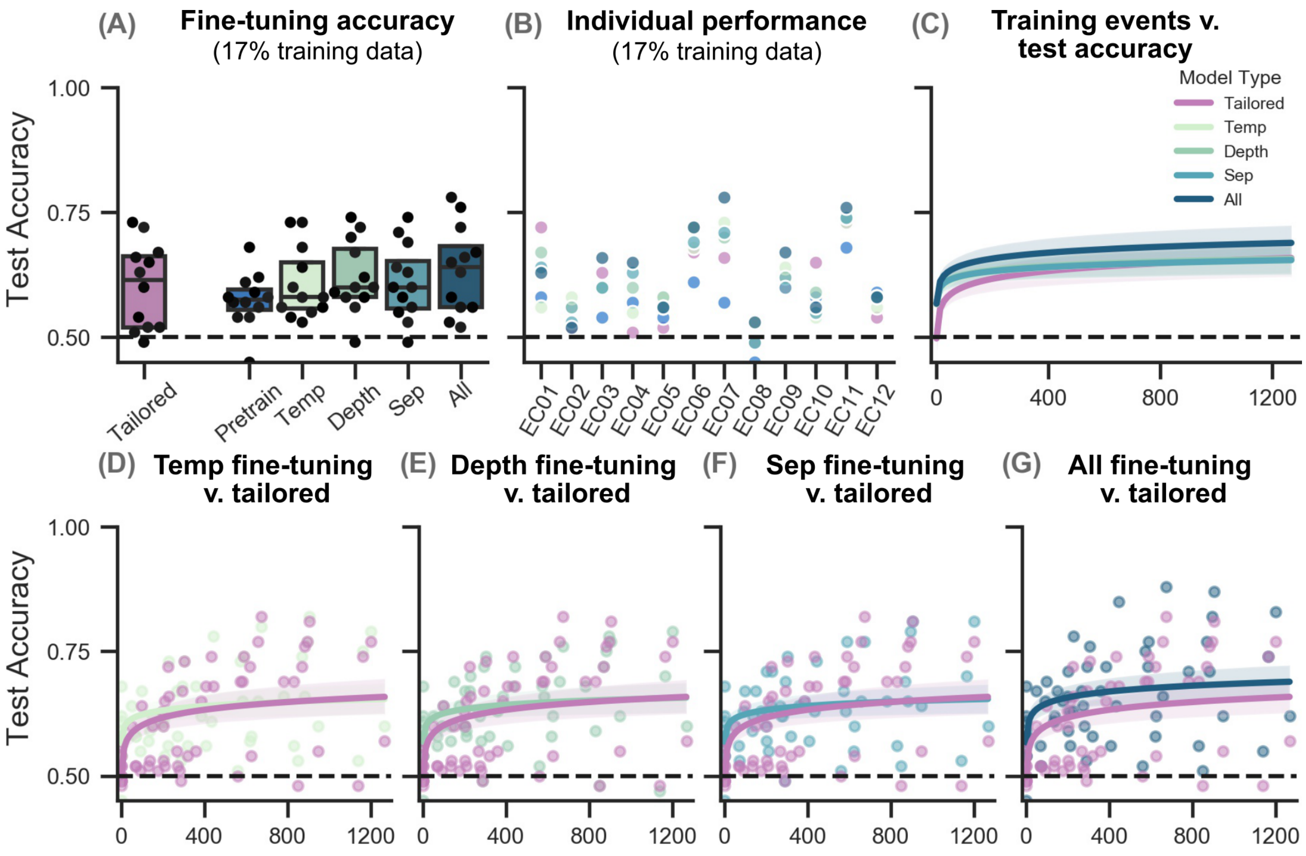


**Figure S4: HTNet decoders transfer from EEG to ECoG participants.** Here, we tested HTNet's ability to transfer from EEG to ECoG data, instead of from ECoG to EEG data as was done in the unseen modality condition. We used relative power with HTNet to transfer between recording modalities, so decoder performance here differs from the tailored and same modality conditions when just power was used. **(A–B)** Fine-tuning all layers improves upon pretrained test accuracy the most compared to the other fine-tuning approaches. **(C)** Fine-tuning all layers on ∼50 events results in decoders with accuracies approaching the performance of the best tailored decoders that were trained on hundreds of events. **(D–G)** We also show fine-tuning curves separated by fine-tuning approach, with each dot indicating the test accuracy of a single fold when trained on either 17%, 33%, 50%, or 67% of available data.

| | Temp | Depth | Sep | All |
|---|---|---|---|---|
| Number of parameters | 532 | 5700 | 4940 | 12238 |
| Same Modality | 62.9 (52.5) | 74.8 (72.8) | 51.7 (49.2) | 66.6 (54.3) |
| Unseen Modality (ECoG to EEG) | 15.9 (12.0) | 10.1 (7.5) | 10.4 (6.8) | 10.0 (5.9) |
| Unseen Modality (EEG to ECoG) | 107.3 (153.3) | 43.9 (39.5) | 37.3 (35.3) | 53.4 (48.2) |

**Table S6: Number of parameters and time to fine-tune pretrained HTNet decoders.** The average training time for our four fine-tuning approaches is shown when training on 33% of available events (standard deviation in parentheses). We performed fine-tuning separately on HTNet's temporal (Temp), depthwise (Depth), and separable (Sep) convolutions, along with fine-tuning all trainable layers (All). Despite having 2–23 times as many parameters as the other approaches, fine-tuning all layers doesn't substantially increase training time on average.

| | Temp | Depth | Sep | All |
|---|---|---|---|---|
| Number of parameters | 532 | 5700 | 4940 | 12238 |
| Same Modality | 29.8 (17.5) | 66.2 (41.8) | 54.8 (32.9) | 30.8 (18.4) |
| Unseen Modality (ECoG to EEG) | 83.4 (78.0) | 51.0 (55.6) | 70.8 (71.2) | 25.8 (38.1) |
| Unseen Modality (EEG to ECoG) | 53.0 (66.4) | 28.8 (23.8) | 29.4 (22.0) | 12.0 (11.0) |

**Table S7: Number of training epochs when fine-tuning pretrained HTNet decoders.** The average number of training epochs during HTNet fine-tuning is shown when training on 33% of available events (standard deviation in parentheses). We separately fine-tuned HTNet's temporal (Temp), depthwise (Depth), and separable (Sep) convolutions, along with fine-tuning all trainable layers (All). While fine-tuning individual convolutional layers took a similar number of training epochs, fine-tuning all trainable layers only required about half as many training epochs on average.

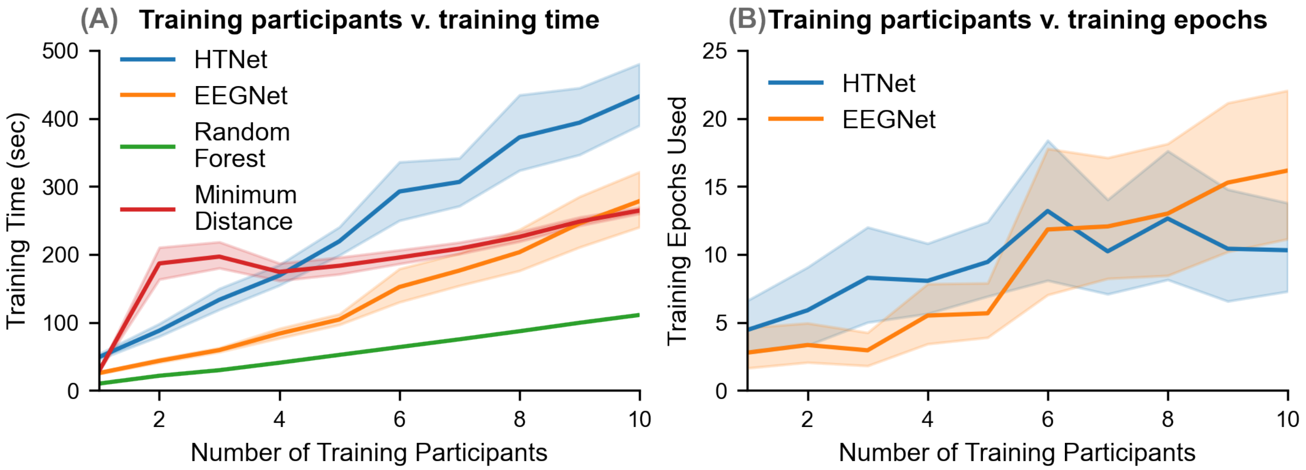


**Figure S5: Decoder training times and epoch numbers for various number of training participants. (A)** As expected, average training time across folds increases with the number of training participants (shading shows 95% confidence interval). While all decoders take under five minutes to train, HTNet's training time is substantially higher than the other decoder types when many training participants are used. **(B)** The number epochs needed for training increases slightly with more training participants, but always remains below 20 epochs on average (shading shows 95% confidence interval).
